# Oral diseases: a 14-year experience of a Chilean institution with a systematic review from eight countries

**DOI:** 10.1101/134056

**Authors:** César Rivera, Carolina Jones-Herrera, Pablo Vargas, Bernardo Venegas, Daniel Droguett

## Abstract

**BACKGROUND:** Retrospective studies to assess the distribution of oral diseases (ODs) are helpful in estimating the prevalence of oral diagnoses in the population, and thus help in preventive and curative services. Prevalence and frequency data for ODs are available from many countries, but information from Chile is scarce.

**METHODS:** This study investigated the frequency of ODs in a Chilean population. For this, we included all patients treated at the University of Talca (UTALCA, Chile) between 2001 and 2014. Patient characteristics were retrieved from medical files. To contextualize our results, we conducted a systematic review (SystRev) using Publish or Perish software (PoP), Google Scholar and MEDLINE/PubMed.

**RESULTS:** One hundred sixty-six ODs were diagnosed, and the most prevalent groups were soft tissue tumours, epithelial pathology and salivary gland pathology. Individually, irritation fibroma, oral lichen planus (OLP) and mucocele were the most common diagnoses. ODs frequently affected unspecified parts of the mouth (including cheek, vestibule and retromolar area), gum, lips, tongue and palate. In the SystRev, the more studied diagnoses were leukoplakia, OLP and recurrent aphthous stomatitis; prevalent lesions included Fordyce’s spots, recurrent aphthous stomatitis and fissured tongue. Chilean patients and SistRev shared almost all ODs.

**CONCLUSION:** The results reflect ODs diagnosed in a specialized service of oral pathology and medicine in Chile and will allow the establishment of preventive/curative policies, adequate health services and dentistry curriculum.

## INTRODUCTION

Oral medicine is a nonsurgical dental discipline that is focused on the diagnosis and management of medical conditions that affect the oral and maxillofacial territories (1). Disorders affecting the oral soft tissues, salivary glands and the oral mucosa are common problems that can affect quality of life (2). Retrospective studies to assess the distribution of oral diseases (ODs) are helpful in estimating the prevalence of oral diagnoses in the population, and thus help in preventive and curative services.

Prevalence and frequency data for ODs are available from many countries, but information from Chile is scarce. The few available studies for the country describe specific diseases, and data are from paediatric and elderly patients.

In Chilean paediatric patients, Zúñiga et al. showed that mucocele was the most commonly found lesion, followed by pyogenic granuloma and irritation fibroma (3). The most common localization for lesions was the lower lip. Previously, two studies coincided in presenting denture-related stomatitis and irritation fibroma as among the most frequent diseases in the Chilean elderly (4, 5). Despite the previous evidence, there are no studies showing the entire range of possible lesions in all age ranges. Hence, with a clinical-only focus, the aim of this research was to determine the frequency of ODs from a Chilean population over a 14-year period. Additionally, we conducted a systematic review to contextualize and compare our results with other countries.

## MATERIALS AND METHODS

### Sample

All patients treated between 2001 and 2014 at the University of Talca were included in the study. Records for 1,009 patients were retrieved from the university archives. The primary inclusion criterion was the availability of clinical and histopathological diagnosis (both). Finally, we conducted a retrospective study with a sample of 1,000 patients (clinicopathological characteristics are presented in Table 1). This study was performed in accordance with the Helsinki Declaration and was approved by the Ethics Review Boards of the University of Talca (2014-027-DD).

**Table 1.**
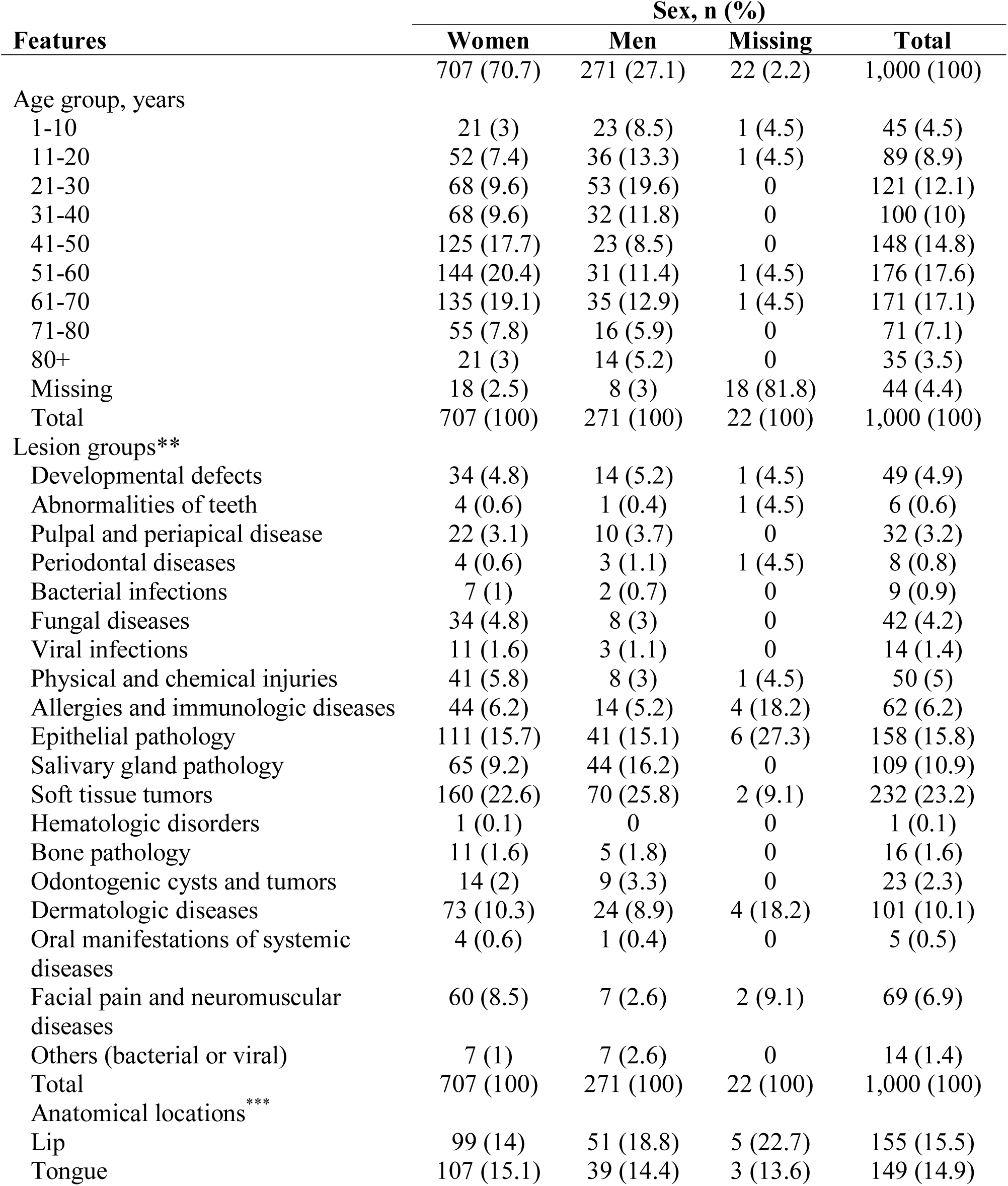
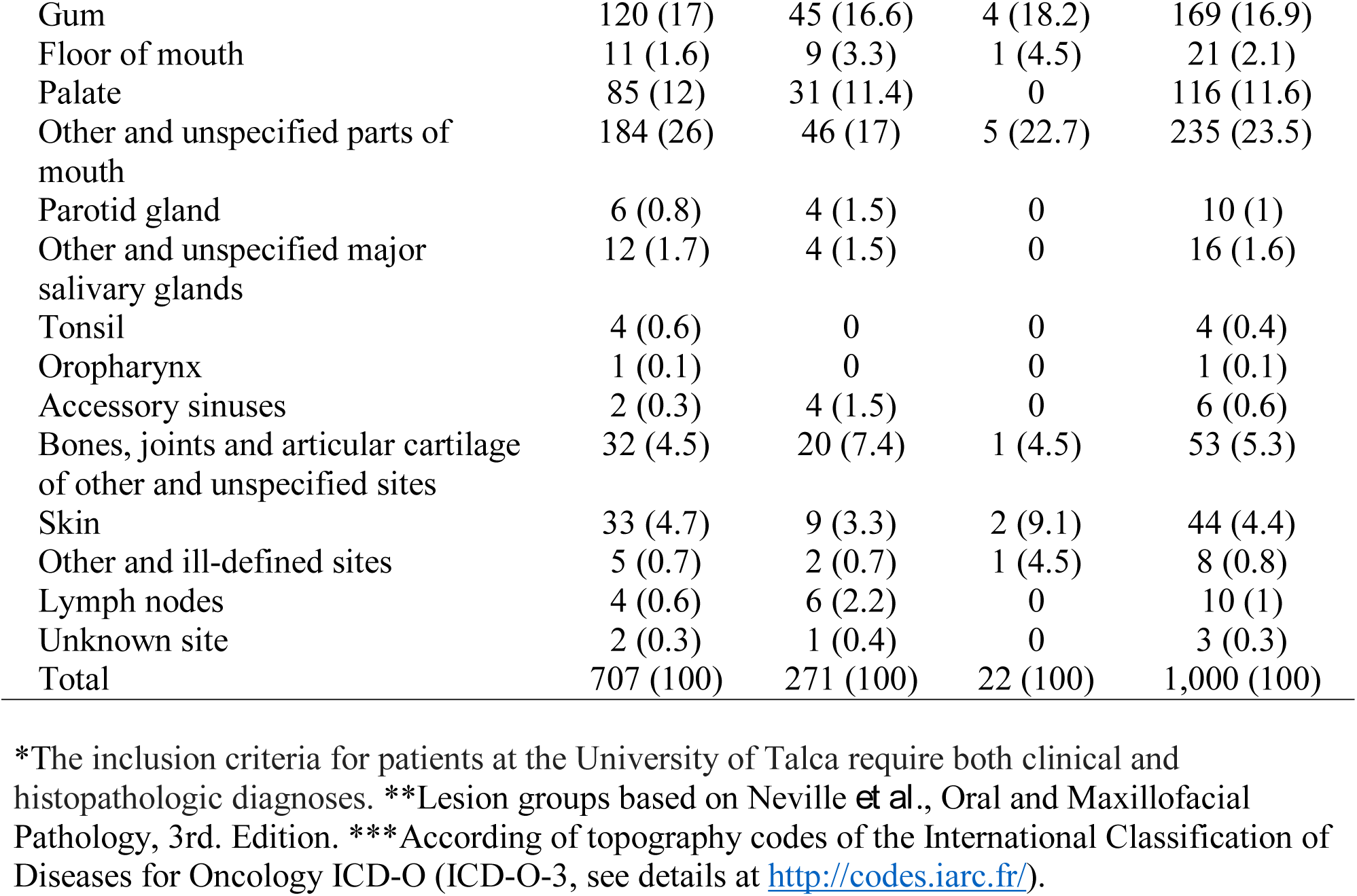
Characteristics of patients* with oral diseases treated at the oral medicine service, University of Talca School of Dentistry, Chile (n=1,000).

### Data collection

ODs were classified according to the textbook of Neville et al. Anatomical sites were reported, according to the International Classification of Diseases for Oncology (ICD-O-3, see details at http://codes.iarc.fr/). The results were informed following the Strengthening the Reporting of Observational Studies in Epidemiology (STROBE) statement.

### Systematic review

To contextualize our results, published articles were identified through a SistRev search with PoP software (http://www.harzing.com/resources/publish-or-perish/windows) using the search operators: “oral lesions” (all the words) and “prevalence frequency” (any of the words), using “title words only” (query date: 2016-04-03). Duplicate articles were eliminated using CleanPoP (http://cleanpop.ifris.net/guide.html). The inclusion criteria were as follows: i) English-language articles, ii) patients with ODs, iii) ODs without specific associations (i.e., HIV, age) and iii) journal available in MEDLINE/PubMed. The exclusion criteria were as follows: i) non-primary content, ii) articles without full access and iii) unclear reports. The results were informed following the Preferred Reporting Items for Systematic Reviews and Meta-Analyses (PRISMA) statement. To study coincidences and logical relations between our sample and the SistRev, we plotted a Venn diagram using the InteractiVenn tool (http://www.interactivenn.net).

### Statistical analysis

The data were analysed with descriptive statistics (frequency and percent) using Microsoft Excel 2013 (Microsoft Corporation, Seattle, USA) and SPSS statistical package 17 for Windows (IBM, Chicago, USA). Missing data were coded under “missing” category.

## RESULTS

### Characteristics of study participants

Publicly available resources (data set) can be accessed at DOI: <u>10.5281/zenodo.164948</u>. Females represented 70.7% of the sample. The average age at diagnosis was 46.5 years (range, 4 to 97 years old), and most patients were in their forties to seventies. There were 166 oral conditions found. The most prevalent groups were soft tissue tumours (232, 23.2%), epithelial pathology (158, 15.8%) and salivary gland pathology (109, 10.9%). Table 1 shows the distribution of patients according to gender, age, lesion groups and anatomical sites. To review underlying diseases, medicines, smoking and drinking habits, please review the data set at DOI: <u>10.5281/zenodo.164948</u>.

### Top ten oral diseases

Table 2 listed the “top ten” ODs (for complete information, see details at DOI: <u>10.5281/zenodo.164948</u>). Individually, irritation fibroma (102 cases, 10.2%), oral lichen planus (58, 5.8%) and mucocele (54, 5.4%) were the most frequent diseases. Oral squamous cell carcinoma represented 0.6% of the sample (only 6 patients). In women, oral lesions were mostly represented by irritation fibroma (102, 10.2%), oral lichen planus (41, 5.8%) and recurrent aphthous stomatitis (30, 4.2%). In men, mucocele (28, 10.3%), irritation fibroma (25, 9.2%) and pyogenic granuloma (17, 6.3%) were the most common oral diseases.

**Table 2.** Top ten oral diseases* in Chilean patients treated at the oral medicine service, University of Talca School of Dentistry.

| Diagnosis | Ranking (n, %) |  |  |  |  |  | Main |  |
| --- | --- | --- | --- | --- | --- | --- | --- | --- |
|  |  | Global |  | Women |  | Men | Age (y) | Anatomic** |
| Irritation fibroma | 1 | (102, 10.2) | 1 | (77, 10.9) | 2 | (25, 9.2) | 51-60 | Cheek mucosa |
| Lichen planus | 2 | (58, 5.8) | 2 | (41, 5.8) | 4 | (14, 5.2) | 51-60 | Cheek mucosa |
| Mucocele | 3 | (54, 5.4) | 5 | (26, 3.7) | 1 | (28, 10.3) | 11-20 | Mucosa of lower lip |
| Burning mouth syndrome | 4 | (45, 4.5) | 2 | (41, 5.8) | 11 | (2, 0.7) | 51-60 | Mouth, NOS |
| Hemangioma | 5 | (41, 4.1) | 4 | (29, 4.1) | 5 | (10, 3.7) | 61-70 | Mucosa of upper lip |
| Recurrent aphthous stomatitis | 6 | (41, 4.1) | 3 | (30, 4.2) | 6 | (9, 3.3) | 21-30 | Cheek mucosa |
| Papilloma | 6 | (38, 3.8) | 6 | (24, 3.4) | 4 | (14, 5.2) | 31-40<br>51-60 | Mucosa of lower lip<br>Palate |
| Pyogenic granuloma | 7 | (32, 3.2) | 9 | (15, 2.1) | 3 | (17, 6.3) | 21-30 | Gum, NOS |
| Frictional hyperkeratosis | 8 | (24, 2.4) | 7 | (18, 2.5) | 8 | (5, 1.8) | 51-60 | Cheek mucosa |
| Melanin pigmentation | 9 | (20, 2) | 8 | (16, 2.3) | 9 | (4, 1.5) | 61-70 | Palate, NOS |
| Nevus | 9 | (20, 2) | 7 | (18, 2.5) | 11 | (2, 0.7) | 21-30 | Cheek mucosa |
| Epulis fissuratum | 10 | (19, 1.9) | 10 | (14, 2) | 8 | (5, 1.8) | 61-70 | Palate, NOS |

\*Global top ten ranking was determined by the number of patients who had the diagnosis.
\*\*According to topography codes of the International Classification of Diseases for Oncology ICD-O (ICD-O-3, see details at http://codes.iarc.fr/). NOS, not otherwise specified. N°10 in the men ranking is shared by 7 diagnoses (3, 1.1% each one).

### Oral diseases according anatomical sites

The conditions frequently affected “unspecified parts of mouth” (including cheek, vestibule and retromolar area, 235 cases, 23.5%), gums (169, 16.9%), lips (155, 15.5%), tongue (149, 14.9%) and palate (116, 11.6%). Table 3 shows diseases with a frequency greater than or equal to five, for each anatomical site. Irritation fibroma commonly affects the cheek mucosa (40, 39.2%) and the mucosa of the lower lip (11, 10.8%). Oral lichen planus (OLP) was mainly found on the cheek mucosa (31, 53.4%) and the tongue (tongue-NOS 8, border of the tongue 5, 22.4%). Mucocele was principally diagnosed on the mucosa of the lower lip (41, 75.9%) and the ventral surface of the tongue (5, 9.3%).

**Table 3.**
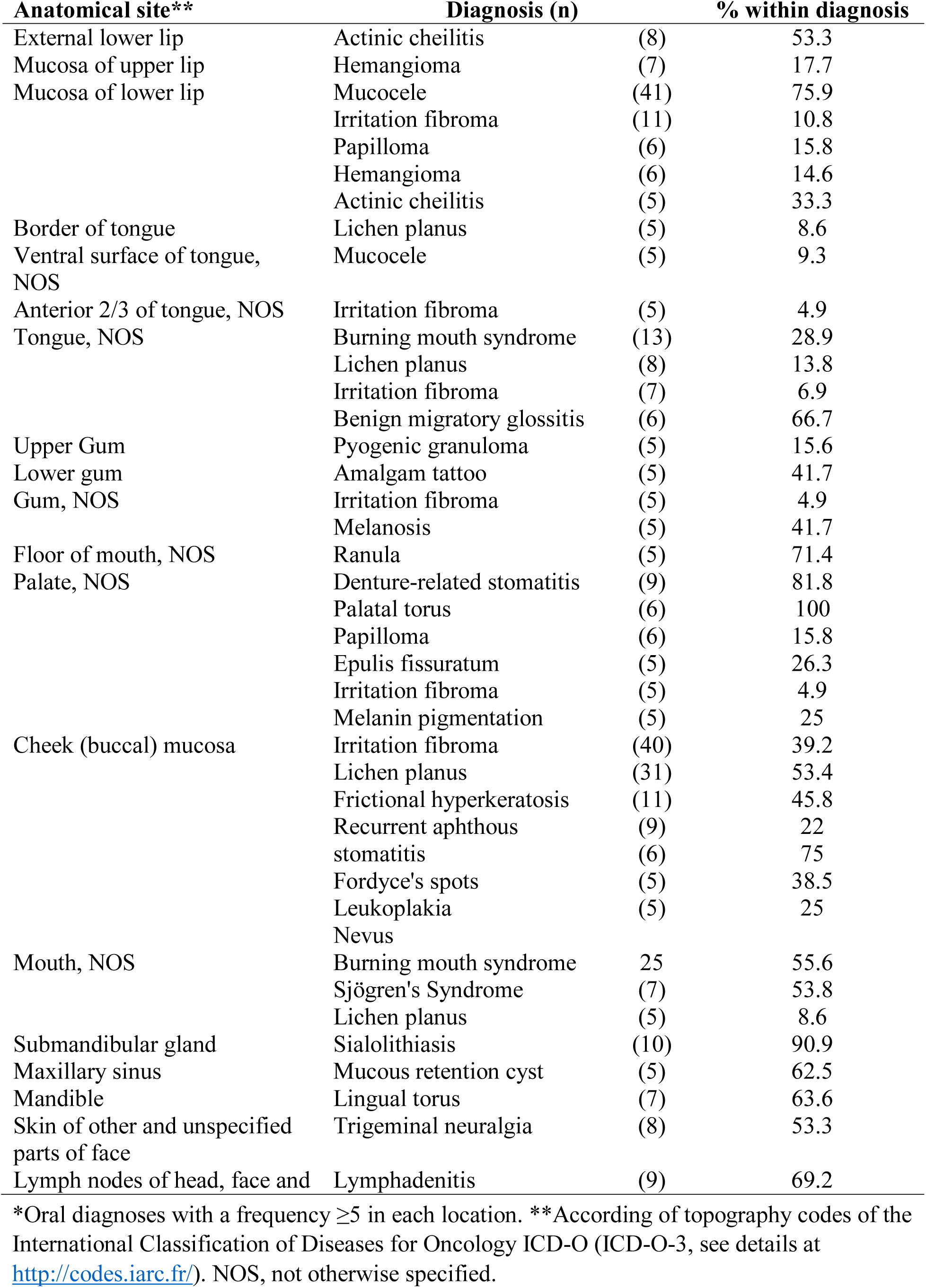
Oral diseases in Chilean patients treated at the oral medicine service, University of Talca School of Dentistry, according anatomical site.

### Consolidation: distribution of oral diseases in our sample

To gain a better understanding of how ODs behaved in our sample, we created a heat map (Figure 1). The map includes ODs with a frequency greater than or equal to ten in our complete sample, a global ranking of lesions (irritation fibroma to radicular cyst), age ranges (1-10 to 80+ years) and anatomical sites (lip to another). For each line, higher numbers represent intense and saturated colours.

**Fig. 1.**
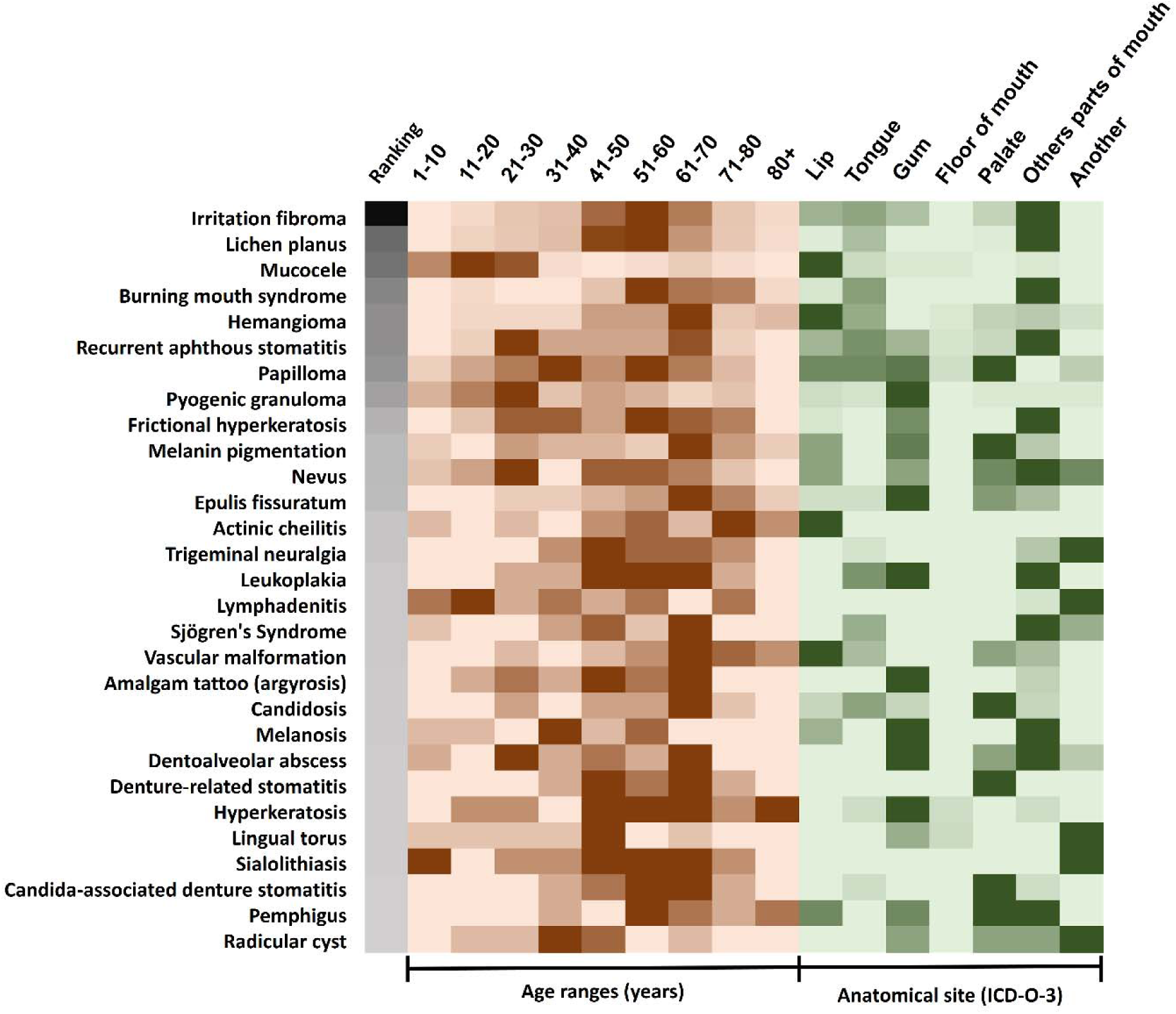
Overview of oral diseases with a frequency ≥10 in a sample of 1,000 Chilean patients. The heat map combines the most frequent ODs (irritation fibroma to radicular cyst), rates by global ranking, age ranges (in years) and anatomical sites (according ICD-O-3). Higher numbers (frequency) represent intense and saturated colours of these lines. In anatomical sites, other and unspecified parts of the mouth include cheek, vestibule and retromolar area (ICD-O-3 C02 code). Another category represents the remaining anatomical sites. In our sample (originated from a specific specialty) some lesions appear to be magnified. For example, oral nevi and trigeminal neuralgia are more common than denture-related stomatitis and radicular cyst. This is explained because denture-related stomatitis and radicular cyst are resolved at the clinic of oral rehabilitation. periodontics and endodontics.

### Systematic review

Figure 2 shows the PRISMA diagram. This strategy identified 325 suitable results, from which 279 were excluded by title review during the screening. Twenty-six were excluded because they are not indexed at MEDLINE/PubMed, and 7 on account of unclear results (or without online access). Full text articles were obtained for 13 studies (6-18), all cross-sectional. See details at DOI: <u>10.5281/zenodo.164948</u>.

**Fig. 2.**
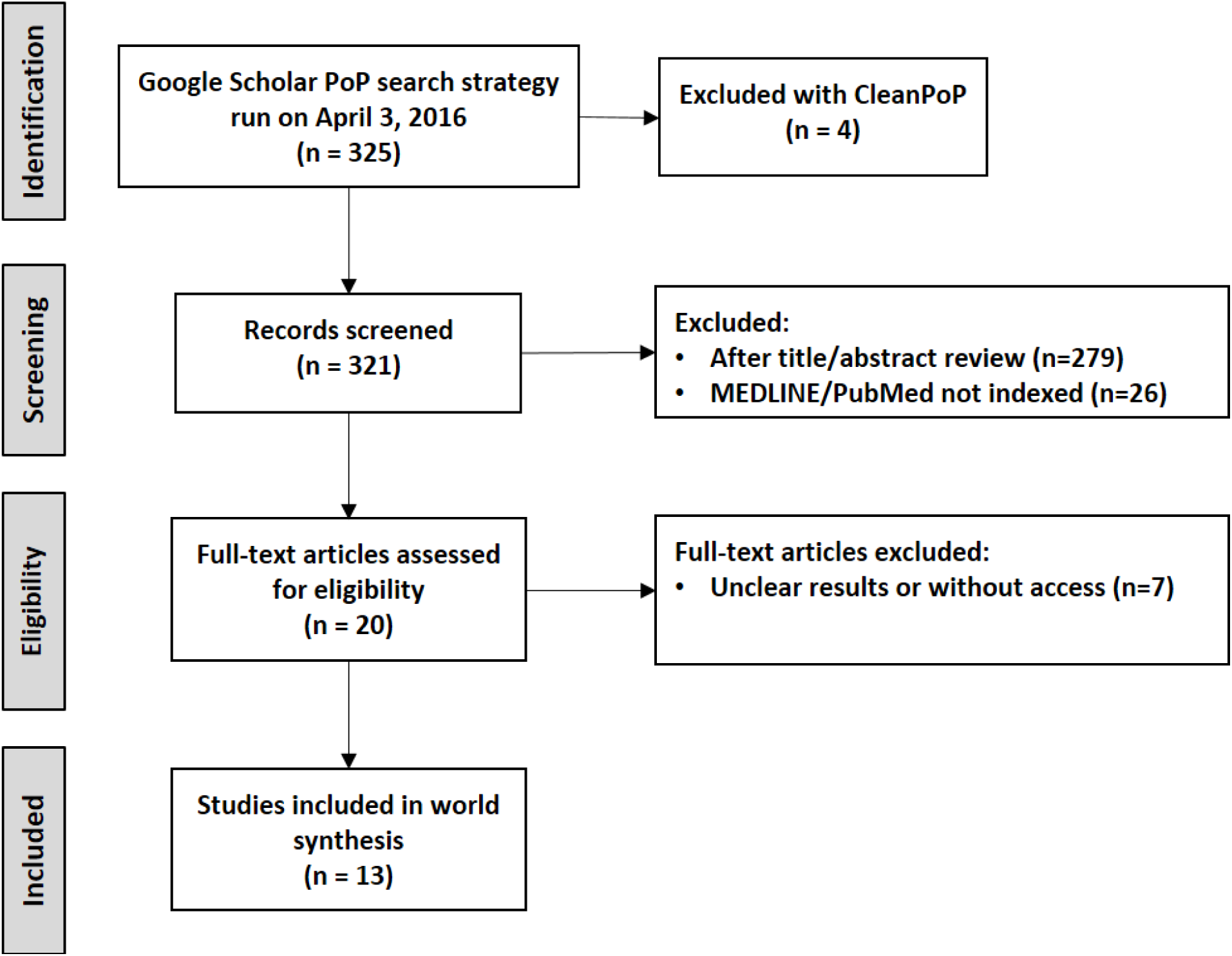
PRISMA diagram of studies searched and selected. We searched the Google Scholar data source using Publish or Perish (PoP) software, based on the terms: “oral lesions” (all the words) and “prevalence frequency” (any of the words), using “title words only”. Duplicate articles were eliminated using CleanPoP.

### Oral diseases around the world

The selected studies were screened, and specific study characteristics were recorded. These parameters are summarized in Table 4. Included studies were conducted in India, Turkey, Saudi Arabia, Slovenia, Cambodia, Brazil, Kuwait, and China. A variable number of patients were reported, ranging from 223 to 24,422 patients. N, age ranges, lesions prevalence percent, and top ODs were extracted.

**Table 4.**
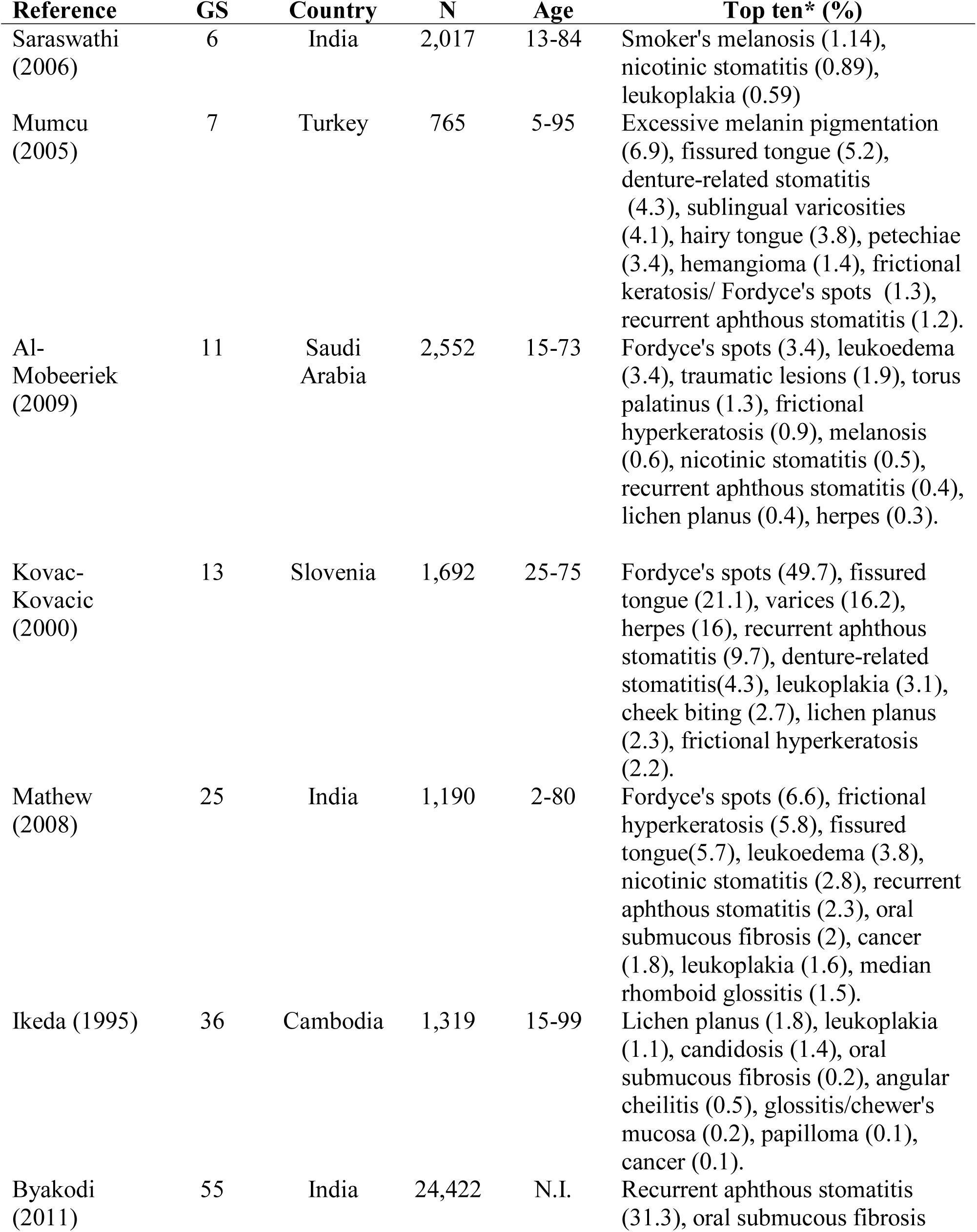
Main characteristics and results of the studies included in the systematic review.

As a synthesis, reported diagnoses were: leukoplakia (8 of 13 articles), oral lichen planus (7/13), recurrent aphthous stomatitis (7/13), Fordyce’s spots (6/13), nicotine stomatitis (6/13), frictional hyperkeratosis (5/13), herpes (5/13), candidiasis (4/13), fissured tongue (4/13), oral submucous fibrosis (5/13) and traumatic ulcers (5/13) (Figure 3A). Considering all studies, Fordyce’s spots, recurrent aphthous stomatitis and fissured tongue were the most frequent ODs.

**Fig. 3.**
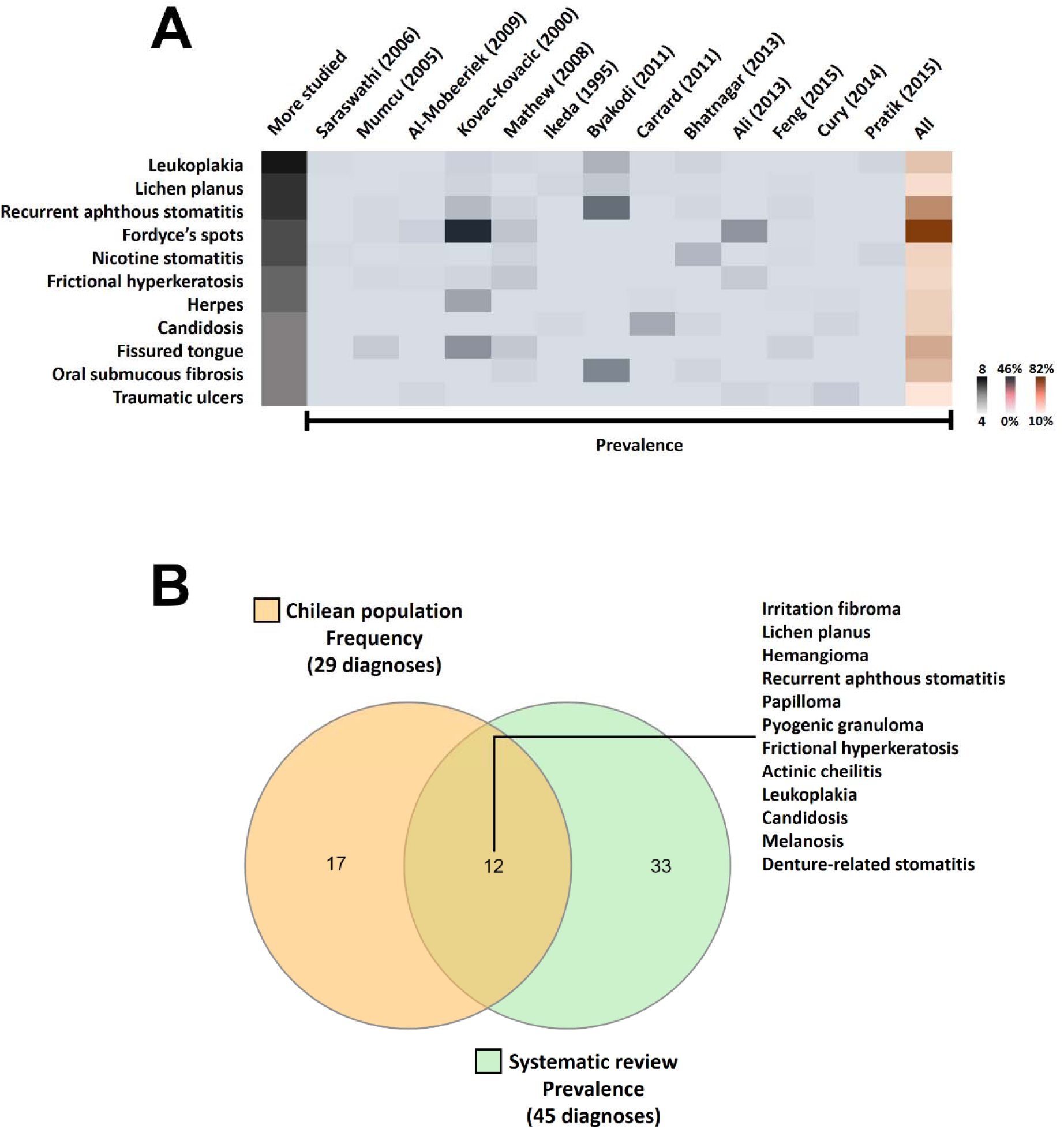
Relevant diagnoses. Top five ranking for more studied lesions in the systematic review (3A). ODs reported in ≥4 articles. The heat map combines the most frequent lesions (leukoplakia to traumatic ulcers), rates by number of articles (grey bar) and prevalence across studies (ordered by Google Scholar rank). Higher numbers represent intense and saturated colours (extreme values on the right-hand side). Venn diagram for the most frequent and prevalent diagnosis (3B). Comparison of rank lists for the current study (our sample), and systematic review (studies around the world). Twelve of the 17 diagnoses (70.6%) in our top ten ranking were shared with selected studies.

### Relation between Chilean diagnoses and systematic review

To understand the similarity between our sample and SysRev articles, we plotted a Venn diagram using the InteractiVenn tool (Figure 3B). Twelve of 17 diagnoses (70.6%) were shared.

## DISCUSSION

Understanding the distribution of ODs is essential to promote prevention, diagnosis, prompt treatment, and the provision of appropriate oral pathology and medicine services. Few reports are available from Chile. To our knowledge, the only ones are in the English-language literature related to the elderly (4, 5) and paediatric populations (3); therefore, it is difficult to make comparisons. To the best of our knowledge, this is the first report describing ODs across all age ranges in our country.

In this study, women had the most frequencies. This outcome may represent a source of potential bias or imprecision because presumably women are more considerate of their oral health than men. In our sample, the conditions frequently affected active mouth anatomical sites during occlusal forces; topographical areas with a role in the mastication efficiency. This could explain some key results (why two of the three most common lesions are trauma-associated); irritation fibroma (or traumatic fibroma) and mucocele.

Irritation fibromas (fibrous hyperplasia) were the most common type of oral disease identified in this research. They are very common hyperplastic lesions of the oral mucosa. These lesions were found more frequently in this study, on the cheek mucosa and lips, findings consistent with classical studies (19, 20). Clinically, irritation fibromas appear either as pedunculated or sessile growths, on any surface of the mucous membrane. The majority, are small lesions that do not have malignant potential, and recurrences mostly result from a failure to eliminate the chronic irritation.

The second most common diagnosis in this study was oral lichen planus (OLP), a relatively common disease of long duration affecting 1-2% of the population, and it has been reported to have an increased potential for malignant transformation to oral cancer. OLP occurs in men and women, especially adults in the 4th–6th decade of life, which is in line with our findings. The red, inflamed lesions and open sores of OLP can cause a burning sensation or pain. The white, lacy patches may not cause discomfort, when they appear on the cheek mucosa, but may be painful when they involve the tongue (21). In our study, OLP affected mainly the cheek mucosa and tongue.

After OLP, mucocele was the next most common lesion. It is a benign pseudo-cystic local lesion (extravasation) that is asymptomatic, usually containing saliva in its interior, and caused by the disruption of the minor salivary gland ducts, or the presence of sialolith inside the ducts. The aetiology of mucocele is trauma-related and the lower lip is the most common area of occurrence of this lesion (22), which is commonly found in children and young adults. These findings are entirely consistent with our results since mucocele occurs on the mucosa of the lower lip in patients between 1 and 30 years old.

To contextualize our results, we conducted a systematic review using Google Scholar (GS) and PoP software. GS displays considerable stability over time and might provide a less-biased comparison across disciplines than the Web of Science (23). PoP retrieves and analyses GS scholar citations and presents a wide range of citation metrics (for example H-index) in a user-friendly format.

Table 4 provides a summary of studies assessing the prevalence of ODs in various populations around the world. It is clear that most lesions are benign, but few have malignant transformation potential, such as leukoplakia, OLP and oral submucous fibrosis (present in the most informed diseases across studies).

In the SysRev Fordyce’s spots, recurrent aphthous stomatitis and fissured tongue were the most frequent ODs. Fordyce’s spots are yellowish, ectopic sebaceous glands. Most of the individuals who develop the granules are males. This condition is entirely benign and does not require any further intervention (24). In our study, Fordyce’s spots were found more frequently among males, and in the 31- to 40-years-old age group.

Recurrent aphthous stomatitis (RAS, or aphthae, aphthous stomatitis), was the second most common diagnosis in the SysRev. RAS is the most common ulcerative disease of the oral mucosa (10-20% of the population), presenting as painful, round, shallow ulcers with a well-defined erythematous margin and a yellowish-grey pseudomembranous centre (25). Diverse factors, including genetic predisposition, immunological problems, viral and/or bacterial infections, allergies, vitamin and microelement deficiencies, diseases, hormonal imbalance, mechanical injuries, and stress, have been suggested to trigger, or to be associated with RAS (26). In our sample, RAS ranks in sixth position.

The third most common diagnosis in the SysRev was fissured tongue. Fissured tongue is relatively common and affects 2% to 5% of the overall population. Deep grooves and fissures on the dorsal surface of the tongue can be seen in children and adults; however, severity and prevalence increases with age (27). This condition does not appear within our most frequent ODs (n°21 in the ranking).

Considering the most common diagnoses in the SysRev (top ten), a large number of our ODs found coincidences (70.6% in Venn diagram); good agreement, considering our study design and sample size.

Although this study provides helpful information, there are several weaknesses. First, the patients treated by the oral pathology and medicine service at The School of Dentistry of University of Talca may not be able to represent ‘an urban population of Chile’. Second, compared to the urban population of Chile, the sample size of 1,000 is also too small. Furthermore, the detection of the lesions was dependent on the knowledge and recognition of diagnoses by multiple examiners in the oral pathology and medicine service, without calibration. Another weakness is that the oral mucosal lesions were diagnosed after only a single examination of each patient, possibly underestimating the frequency of recurrent and chronic alterations. Finally, another problem is trying to compare oral abnormalities diagnosed in a specific specialty setting to previous studies done on more representative general populations.

In Chile, prevalence studies are needed to determine the actual prevalence of ODs in the population. Despite the limitations, the results of this research should be useful in providing more information about oral diseases in the Chilean population and in Latin Americans. In conclusion, our results provide important information about the frequency of ODs in an urban population of Chile. The information presented in this study adds to our understanding about the common oral diseases occurring in the general population and will allow the establishment of preventive policies, adequate oral medicine clinical services and emphasis on dentistry curricula.

## ACKNOWLEDGES

CR is a beneficiary of Chile’s National Commission for Scientific and Technological Research (CONICYT) Becas-Chile scholarship for Ph.D. students, No. 8540/2014. DD is a beneficiary of CONICYT national scholarship for Ph.D. students, No. 21120391/2012. CR & DD are corresponding authors. The authors thank the Maule region patients for giving us the opportunity to undertake this research in their community and Professors Bernardita Fuentes, Marcelo Sánchez, Sonia Vázquez and Wendy Donoso (oral pathology team) for professional and logistical support.

